# Somatosensory responses to nothing: an MEG study of expectations during omission of tactile stimulations

**DOI:** 10.1101/336479

**Authors:** Lau M. Andersen, Daniel Lundqvist

**Affiliations:** NatMEG, Department of Clinical Neuroscience, Karolinska Institutet, Nobels väg 9, 171 77 Stockholm, Sweden

**Keywords:** expectations, somatosensory processing, magnetoencephalography, time-keeping, mismatch responses, cerebellum

## Abstract

The brain builds up expectations to future events based on the patterns of past events. This function has been studied extensively in the auditory and visual domains using various oddball paradigms, but only little exploration of this phenomenon has been done in the somatosensory domain. In this study, we explore how expectations of somatosensory stimulations are established and expressed in neural activity as measured with magnetoencephalography. Using tactile stimulations to the index finger, we compared conditions with *actual stimulation* to conditions with *omitted stimulations*, both of which were either *expected* or *unexpected*.

Our results show that when a stimulation is expected but omitted, a time-locked response occurs ∼135 ms subsequent to the expected stimulation. This somatosensory response to “nothing” was source localized to the secondary somatosensory cortex and to the insula. This provides novel evidence of the capability of the brain of millisecond time-keeping of somatosensory patterns across intervals of 3000 ms.

Our results also show that when stimuli are repeated and expectations are established, there is associated activity in the theta and beta bands. These theta and beta band expressions of expectation were localized to the primary somatosensory area, inferior parietal cortex and cerebellum. Furthermore, there was gamma band activity in the right insula for the first stimulation after an omission, which indicates the detection of a new stimulation event after an expected pattern has been broken.

Finally, our results show that cerebellum play a crucial role in predicting upcoming stimulation and in predicting when stimulation may begin again.

## Introduction

Conceiving of the brain as not only a passive recipient of stimulation, but also as an active predictor of future stimulation dates back to at least Helmholtz (1867). In support of this notion, a seminal electrophysiological experiment (Näätänen et al., 1978)) demonstrated that the auditory cortex generates a characteristic response to deviant sounds in a sequence of otherwise standard sounds. This response, which manifested as a time-locked increased negativity in the electroencephalogram (EEG) from about 130 ms to about 300 ms after stimulus onset of the deviant stimuli over fronto-central electrodes, was coined the MisMatch Negativity (MMN). The MMN has subsequently been explored in numerous magnetoencephalography (MEG) and EEG studies for auditory pattern deviations in form of frequency, intensity, and duration shifts (Giard et al., 1995) and also in form of phonemic deviations (Näätänen et al., 1997). Demonstrating the involvement of prediction in these responses, an MMN has even been found when the deviant “stimulation” is in the form of a complete omission of sound, but only then if the latency between sounds is briefer than ∼150 ms (Yabe et al., 1997). The neural generators of the auditory MMN have been localized to the primary and secondary auditory cortices (Alho, 1995).

MMNs have also been reported in the visual (Pazo-Alvarez et al., 2003) and somatosensory (Karhu and Tesche, 1999; Tesche and Karhu, 2000) domains. Karhu and Tesche (1999) used MEG to investigate neural responses to trains of median nerve stimulations applied at a 2 Hz rate with random omissions occurring 15 % of the time. However, while including omissions in their stimulation sequences, these authors focussed on the differences between first stimulations after an omission and the remaining stimulations, and thus did not explore time-locked responses to the omissions themselves. In a follow-up study (Tesche and Karhu, 2000) however, the authors reported induced cerebellar activity in the theta and gamma bands after omissions of stimuli.

Most researchers on this topic have studied prediction using tasks where pattern deviations are executed in form of variations in the actual sensory stimulation (e.g. frequency or intensity). For researchers interested in responses to violations of expectations, such stimulation variations offer a very useful approach to map out the precision of the expectations and the sensitivity to violations. However, for those interested in the expectations themselves, the approach contaminates the response of interest, since the neural response to a stimulation event is formed by a combination of exogenous sensory stimulation, endogenous event expectations and a neural response to the updating of that expectation. In our study, we aimed to separate the effects from the exogenous sensory stimulation from the endogenous event expectations.

There has been renewed interest in the brain functions involved in expectations of somatosensory events. Allen et al. (2016) using functional Magnetic Resonance Imaging (fMRI), investigated connective properties between areas of the brain when stimulations unexpectedly shifted from one hand to the other. They found that the thalamus, the insula, the primary somatosensory cortex (SI), middle cingulate cortex (MCC) and the middle frontal gyrus (MFG) all show greater BOLD-responses for deviant stimulations than for expected stimulations. Fardo et al. (2017) conducted an MEG study, also investigating the responses to stimulations unexpectedly shifting from one hand to the other, and furthermore found the inferior parietal cortex (IPC) and the inferior frontal gyrus (IFG) to be involved.

### Purpose and aims

In this study, we aimed to explore how expectations of somatosensory stimulations are established and expressed. For this purpose, we used regular sequences of tactile stimulations at fixed intervals that were irregularly interrupted by omitted stimulations. To accomplish analysis of the neural responses to these events by means of both time-locked and induced analyses, we used a long inter-stimulus interval of 3000 ms. This ISI allowed us to explore expectancy responses related to the periods *before*, *at*, and *after* the point in time where stimulations occurred (or should have occurred, when omitted). The two questions of particular interest – both targeting the expression of expectations – were:

1) Do responses *to stimulations* differ when comparing a condition where (i) an expectation has been built up to (ii) one where an expectation has *not* been built up? (Table 1: *Repeated Stimulation* versus *First Stimulation*).

**Table 1:** Labelling of conditions as to whether stimulations and expectations are present. The labels will be further explicated in the methods section.

|  | Expectation | No Expectation |
| --- | --- | --- |
| Stimulation | (i) *Repeated Stimulation* | (ii) *First Stimulation* |
| No Stimulation | (iii) *Omitted Stimulation* | (iv) *Non-Stimulation* |

2) Do responses to *no stimulations* differ when comparing (iii) a condition where an expectation has been built up with (iv) one where an expectation has *not* been built up? (Table 1: *Omitted Stimulation* versus *Non-Stimulation)*.

For the time-locked responses, we were mainly interested in the first two early components following tactile stimulations: the contralateral SI component after ∼60 ms, and the bilateral secondary somatosensory cortex (SII) component after ∼135 ms (Hari et al., 1984). For the induced responses, we were mainly interested in differential activity between stimulation and omissions in the mu-bands (mu-alpha: ∼8-12 Hz and mu-beta: ∼18-25 Hz). The theta (∼4-7 Hz) and gamma (∼40-100 Hz) bands were also of interest since several studies have shown the involvement of these bands in encoding memories (Nyhus and Curran, 2010; Osipova et al., 2006; Roux and Uhlhaas, 2014; Schack et al., 2002).

In terms of localizing activity related to stimulations and unexpected omissions, we chose to focus on a list of *a priori* areas based on the findings of Allen et al. (2016), Fardo et al. (2017) and Tesche and Karhu (2000). This included the insula, the thalamus, the middle cingulate cortex (MCC), the middle frontal gyrus (MFG), the primary somatosensory cortex (SI), the inferior parietal cortex (IPC), the inferior frontal gyrus (IFG) and the cerebellum. If cerebellum is involved, activity should be seen ipsilaterally (Tesche and Karhu, 1997).

MEG differs in the sensitivity that it has to these areas: especially thalamic activity may be challenging to localize due to its sub-cortical position (Hämäläinen et al., 1993). Insula and MCC are also in parts of the brain where MEG has low sensitivity (Hillebrand and Barnes, 2002), but where localization may still be feasible. SI and MFG, IFG and IPC however are cortical and thus good targets for MEG. Finally, more and more evidence is surfacing for MEG being sensitive to deep sources (e.g. Attal et al., 2012; Coffey et al., 2016; Garrido et al., 2015; Tenney et al., 2013), demonstrating that localization of cerebellar and thalamic activity is feasible.

## Methods

### Participants

Twenty participants volunteered to take part in the experiment (eight males, twelve females, Mean Age: 28.7 y; Minimum Age: 21; Maximum Age: 47). The experiment was approved by the local ethics committee, “Regionala etikprövningsnämnden i Stockholm”, in accordance with the Declaration of Helsinki.

### Stimuli and procedure

Tactile stimulations were generated by using an inflatable membrane (MEG International Services Ltd., Coquitlam, Canada) fastened to the participants’ right index fingertips. The membrane was part of a custom stimulation rig, and was controlled by pneumatic valves (model SYJ712M-SMU-01F-Q, SMC Corporation, Tokyo, Japan) using 1 bar of pressurized air. The experimental paradigm consisted of inflating the membrane with a regular interval of exactly 3000 milliseconds. At pseudo-random intervals, some of the inflations were omitted such that there was a complete absence of stimulation. These omissions always happened at either the fourth, fifth or sixth place in the stimulation sequence; chosen in a counterbalanced manner. An illustration of the sequence is shown in Fig. 1.

**Fig. 1.**
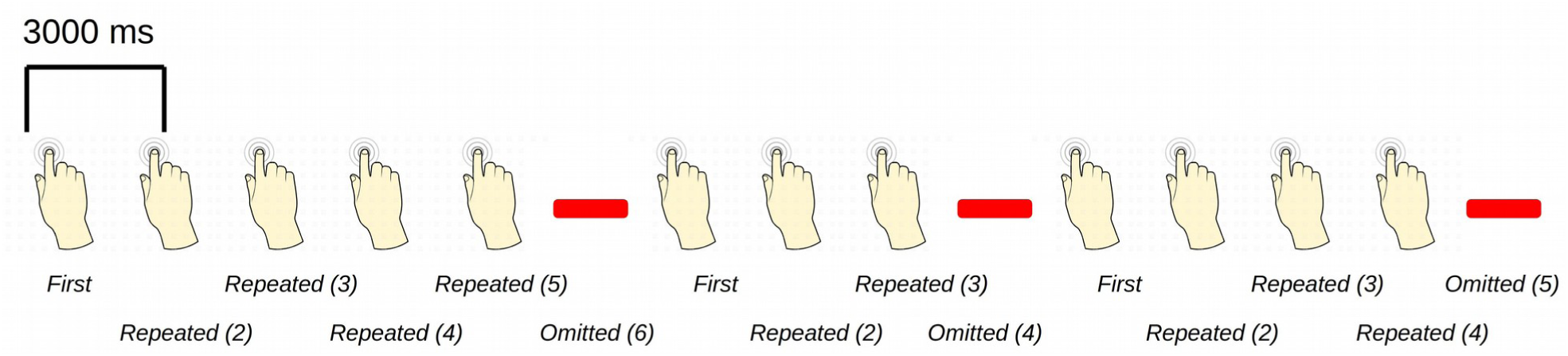
Experimental paradigm: An example sequence of the experimental paradigm is shown. The annotations on the bottom show the coding used throughout for the different events of interest. Stimulations happened at a regular pace, every three seconds. When omissions occurred, there were thus six seconds between two consecutive stimulations.

One thousand trials were administered to each participant, of which six hundred were equally distributed as *First Stimulation, Repeated Stimulation (2)* and *Repeated Stimulation (3).* Two hundred were omissions, close to evenly distributed between *Omitted Stimulation (4), Omitted Stimulation (5)* and *Omitted Stimulation (6)*. The last two hundred were distributed between *Repeated Stimulation (4)* and *Repeated Stimulation (5)*. Periods of fifteen seconds of non-stimulation were interspersed between these sequences to obtain segments of data with repeated omitted stimulation. These occurred approximately every twenty-five trials, always started after an omission, and were cued by a three-second tone. The first three seconds were a *wash-out period,* and also polluted by the tone, and thereafter four trials (3000 ms) of non-stimulation trials were segmented from the remaining twelve seconds of non-stimulation. This resulted in a total of around one-hundred-thirty epochs with no stimulation per subject (*Non-Stimulation*).

During the stimulation procedure, participants were watching a nature programme with sound being fed through sound tubes (model ADU1c, KAR Oy, Helsinki, Finland) into the ears of participants at approximately 65 dB, rendering the tactile stimulation completely inaudible. Participants were instructed to pay full attention to the movie and to pay no attention to the stimulation of their finger, which was held under a table such that it could not be seen. In this way, expectations should be mainly stimulus driven, and thus not cognitively driven or attention driven. Both before and after the administration of the one thousand trials, a period of non-stimulation lasting 3 minutes was recorded. These were cut into segments of 3000 ms, resulting in 120 *No-Task* trials recorded outside the experiment. These were to be used as a common baseline activity for all four conditions (Table 1).

### Preparation of subjects

In preparation for the MEG-measurement each subject had their head shape digitized using a Polhemus FASTRAK. Three fiducial points, the nasion and the left and right pre-auricular points, were digitized along with the positions of four head-position indicator coils (HPI-coils). Furthermore, about 200 extra points digitizing the head shape of each subject were acquired.

### Acquisition of data

Data was recorded by an Elekta Neuromag TRIUX system inside a magnetically shielded room (model Ak3b (Vacuumschmelze GmbH) at a sampling frequency of 1000 Hz and on-line low-pass and high-pass filtered at 330 Hz and 0.1 Hz respectively.

### Processing of MEG data

Two kinds of analyses were done. In the first set of analyses, we extracted responses time-locked to the onset of the actual stimulation (*Repeated* and *First Stimulation*) and to the expected onset of the *Omitted Stimulations*. In the second set of analyses, we extracted induced responses to the same conditions. Both kinds of analyses were also done for *Non-Stimulations.*

#### Time-locked responses

The data were first MaxFiltered (Taulu and Simola, 2006), using temporal Signal Space Separation (tSSS) with a correlation limit of 98 %, movement corrected and line-band filtered (50 Hz). Subsequently the data were low-pass filtered at 70 Hz and then cut into segments of 1200 ms, 200 ms pre-stimulus and 1000 ms post-stimulus. The delay between the digital trigger and the onset of the stimulation was assessed to be 41.0 ms via a separate recording using an accelerometer attached to the tactile membrane. After subtracting the 41.0 ms delay, data was then demeaned using the pre-stimulus period. Segments of data including magnetometer responses greater than 4 pT or gradiometer responses greater than 400 pT/m were rejected. An independent component analysis was done on the segmented data to identify eye blink and heart beat related components. These were subsequently removed from the segmented data. The omissions (occurring at position 4, 5 or 6 in a stimulation sequence) were collapsed into one response category of *Omitted Stimulations* to maximize the signal-to-noise ratio.

#### Induced responses

The data were first MaxFiltered (Taulu and Simola, 2006), using tSSS with a correlation limit of 98 %, movement corrected and line-band filtered (50 Hz). Subsequently, the data were then cut into segments of 3000 ms, 1500 ms pre-stimulus and 1500 ms post-stimulus adjusted with the measured delay of 41.0 ms. Data was demeaned using the whole segment. Then data was cleaned manually by removing segments showing large variance. An independent component analysis was done on the segmented data to identify eye blink, eye movement and heart beat related components. These were subsequently removed from the segmented data. Then, from each segment of data the respective time-locked response of the given condition was subtracted from that segment. This was done to minimize the presence of time-locked responses in the induced responses. Time-frequency representations were calculated from these segments of data according to condition. A Morlet wavelet analysis with 7 cycles was done for frequencies from 1-40 Hz and a multitaper analysis with 5 tapers was done for frequencies from 40-100 Hz. These were done for each time point from 1500 ms pre-stimulus to 1500 ms post-stimulus. Data from gradiometer pairs were then combined by summing the powers from each. Finally, the data were baselined by using the mean power from the *No-Task* trials. To choose which time and frequency ranges to run source reconstruction on, cluster statistics (Maris and Oostenveld, 2007) was done on the differences between *Repeated* and *First Stimulation* and between *Omitted* and *Non-Stimulation*. To this end, separate mass-univariate tests were run on the differences respectively between *Repeated* and *First Stimulation* and between *Omitted* and *Non-Stimulation* with α = 0.05. Individual time-frequency points were considered to be part of a cluster if they were significant on this test *and* they had neighbouring points in space, i.e. sensors, and time-frequency of the same sign, positive/negative, that were also significant. For each positive and negative cluster, the *t*-values were then summed together giving a *T*-value. Subsequently, from a permutation test with 2000 repetitions, labels, e.g. *Repeated* and *First*, were subsequently randomly allocated to each of the conditions, giving a distribution of summed cluster values to compare against, i.e. a distribution of *T*-values. Clusters were considered significant if the likelihood of the *T*-value, or of a more extreme *T*-value, of such a cluster was 0.025 or smaller under the permutation distribution. This controls the false alarm rate for clusters at a level of α = 0.05. Importantly, the multiple comparisons problem is circumvented in this manner. The overall false alarm rate is controlled at a level of α = 0.05, since the null hypothesis, namely that labelling of conditions into, say, *Repeated* and *First Stimulation* is not different than any other (random) labellings of conditions, is only rejected if at least one negative or one positive cluster has a *T*-value that is associated with a *p*-value equal to or smaller than 0.025 under the permutation distribution.

### Source reconstruction

Two different strategies were followed for the source reconstruction of time-locked and induced responses respectively. For the time-locked responses, a distributed solution was found and for the induced responses a beamformer approach was used. See sections below for further details.

#### Time-locked responses

For each subject a set of T1 MR images were acquired using a 3 T GE 750. Based on these images a full segmentation of the head and the brain was done using FreeSurfer (Dale et al., 1999; Fischl et al., 1999a). Based on this segmentation, the boundaries for the skin, skull and brain surfaces were found using the watershed algorithm with the MNE-C software (Gramfort et al., 2013). A source space restricting sources to the cortical sheet was created and a single compartment volume conductor model was set up based on the boundary for the brain surface, also using MNE-C. For each subject the T1 was co-registered to the subject’s head shape with the fiducials and head shape points acquired with the Polhemus FASTRAK. A forward model was then made based on the transformation, the volume conductor, the source space model and the positions of the MEG sensors, (magnetometers and gradiometers). Source time courses were reconstructed using the Minimum-Norm Estimate (MNE) (Hämäläinen and Ilmoniemi, 1994), with depth-weighting (Dale et al., 2000). The noise-estimate necessary for doing the MNE was estimated based on the pre-stimulus activity. The individual source time courses were then morphed onto a common template, the *fsaverage* (Fischl et al., 1999b), from the FreeSurfer software. Grand averages were then done over the morphed data.

#### Induced responses

After applying the transformation done above, we used the FieldTrip software (Oostenveld et al., 2011) to segment the T1-images into the brain, skull and skin for each subject. From the brain segmentation a single compartment volume conductor was created. The source space was the whole brain, and was thus not restricted to the cortical sheet as in the reconstruction of the time-locked responses. A beamformer approach was used (Dynamic Imaging of Coherent Sources, DICS: Gross et al., 2001). Based on the identified components in the time-frequency representations, trial segments were cropped into time windows containing the component. From these time windows, the Fourier transforms for the frequency of interest for the *Repeated* and *First Stimulation* and the *Omitted* and *Non-Stimulation* were done. Furthermore, a Fourier transform was also done for the *No-Task* trials collected outside the experiment, to be used as a contrasting condition for all conditions (Table 1). Finally, Fourier transforms were done for the combinations of each of the conditions and the *No-Task* trials outside the experiment. Based on these Fourier transforms, sources underlying the induced responses were reconstructed with the usage of a common spatial filter estimated from the combinations of the conditions and the *No-Task* trials outside the experiment. The co-registered source space for each subject was warped onto the standard Colin-27 brain (Holmes et al., 1998). The forward model for each subject was based on the warped source space, the positions of the gradiometers and the volume conductor. Contrasts between beamformer reconstructed activity for each of the conditions (*First, Repeated, Omitted* and *Non-Stimulations*) and *No-Task* trials outside the experiment were calculated. These revealed which regions generated the induced responses.

## Results

Note that for all results *Repeated Stimulation* is the second stimulation (Fig. 1 and Table 1), i.e. the one following *First Stimulation*, since this is where the expectation of another stimulation can be confirmed. The reserved digital object identifier for the data repository, where data for this experiment can be freely downloaded is: 10.5281/zenodo.998518.

### Time-locked responses

Grand averages were calculated across all participants separately for each of the conditions. The sensor space analyses were mostly done for quality inspection of the data (Fig. 2) and will therefore not contain any statistical analyses, as these will be done in the source space.

**Fig. 2.**
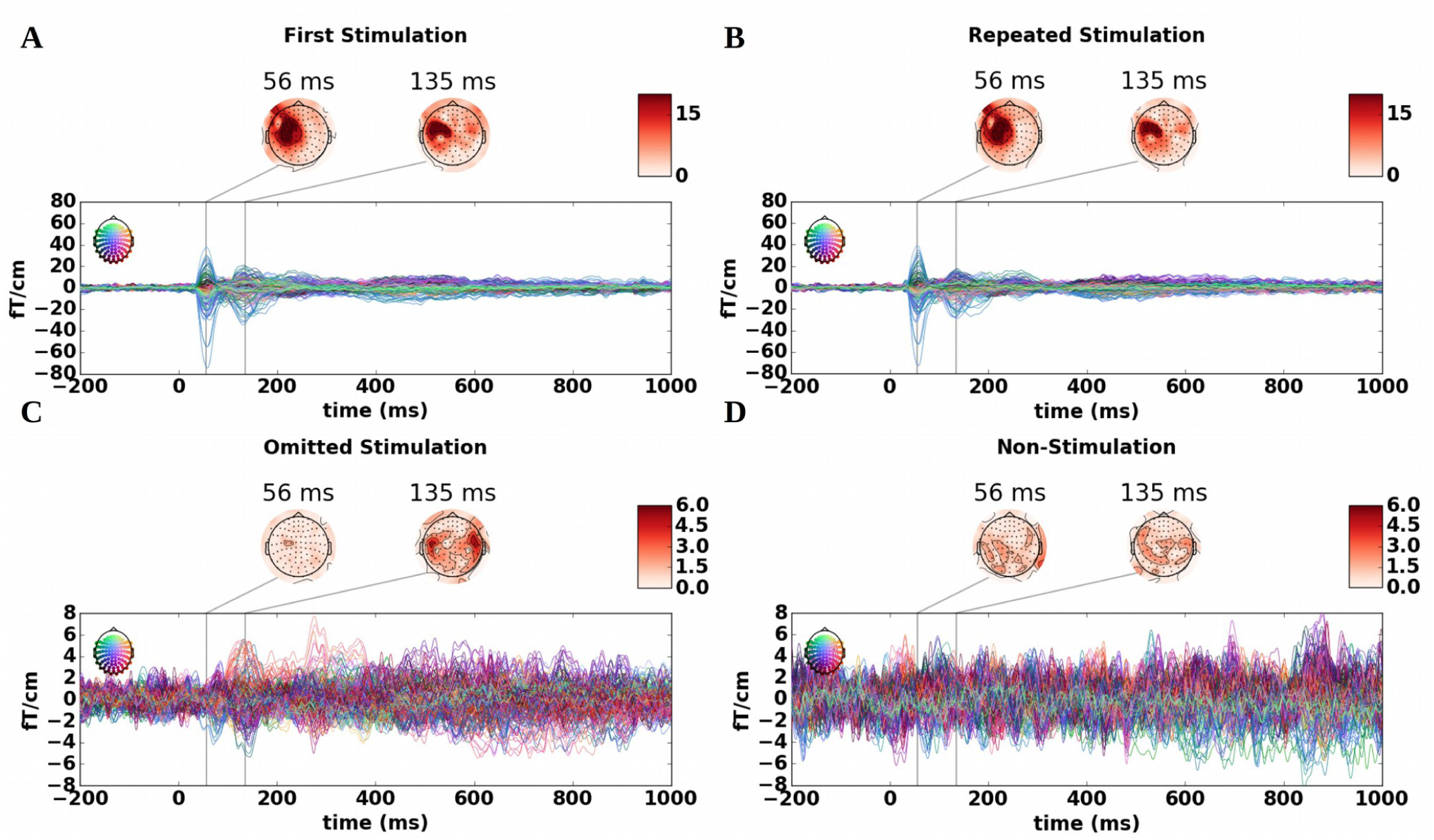
Timelocked responses: Butterfly plots of the time-locked responses for the 204 gradiometers with accompanying flattened topographical plots with root mean square values of the gradiometer pairs. Coloured heads in the top-left corners of the butterfly plots indicate from which sensors data are drawn. **A)** contralateral peak for *First Stimulation* after 56 ms (SI) and bilateral peak after 135 ms (SII). **B)** similar to **A**, but for *Repeated Stimulation*. **C)** a bilateral peak after 135 ms with SII topography. **D)** For the *Non-Stimulations*, no clear components were found. Note the different scales in the top row and the bottom row.

*Repeated* and *First Stimulation* were compared to one another, and so were *Omitted* to *Non-Stimulation* (Table 1). The results showed that *Repeated* and *First Stimulation* were very similar with the first component occurring contralaterally to the stimulated hand at 56 ms over the somatosensory cortex (SI-component), which was followed by a second bilateral component at 135 ms over the secondary somatosensory cortices (SII-component) (Fig. 2AB). The *Omitted Stimulation* lacked the initial SI-component observed for stimulation conditions at 56 ms, but showed an SII-component at 135 ms, thus matching the timing of the second component for both *Repeated* and *First Stimulation* (Fig. 2C). *Non-Stimulation* did not show any systematic components at any of these time points (Fig. 2D).

The SI and SII components shown in Figs. 2A and 2B were expected after tactile stimulation, because the responses in the somatosensory cortex are known to be unfolding at a very reliable and precise pace following sensory stimulations (Hari and Forss, 1999). The time-locked responses to *Omitted Stimulation* were however more unexpected (Fig. 2C).

#### Source reconstruction of time-locked responses

Using Minimum-Norm Estimates (as described in the methods) we reconstructed the sources for the two responses (56 ms (SI) and 135 ms (SII); see Fig. 2). To test the statistical significance of these responses, we performed permutation tests (Maris and Oostenveld, 2007) with a threshold of α = 0.05 for the initial mass-univariate test and a subsequent cluster threshold of α = 0.025 (Fig. 3). The data going into the test was the difference between source reconstructed time courses morphed to *fsaverage* with a 10 ms interval around the peak (i.e. 46-66 ms and 125-145 ms) intervals were tested.

**Fig. 3.**
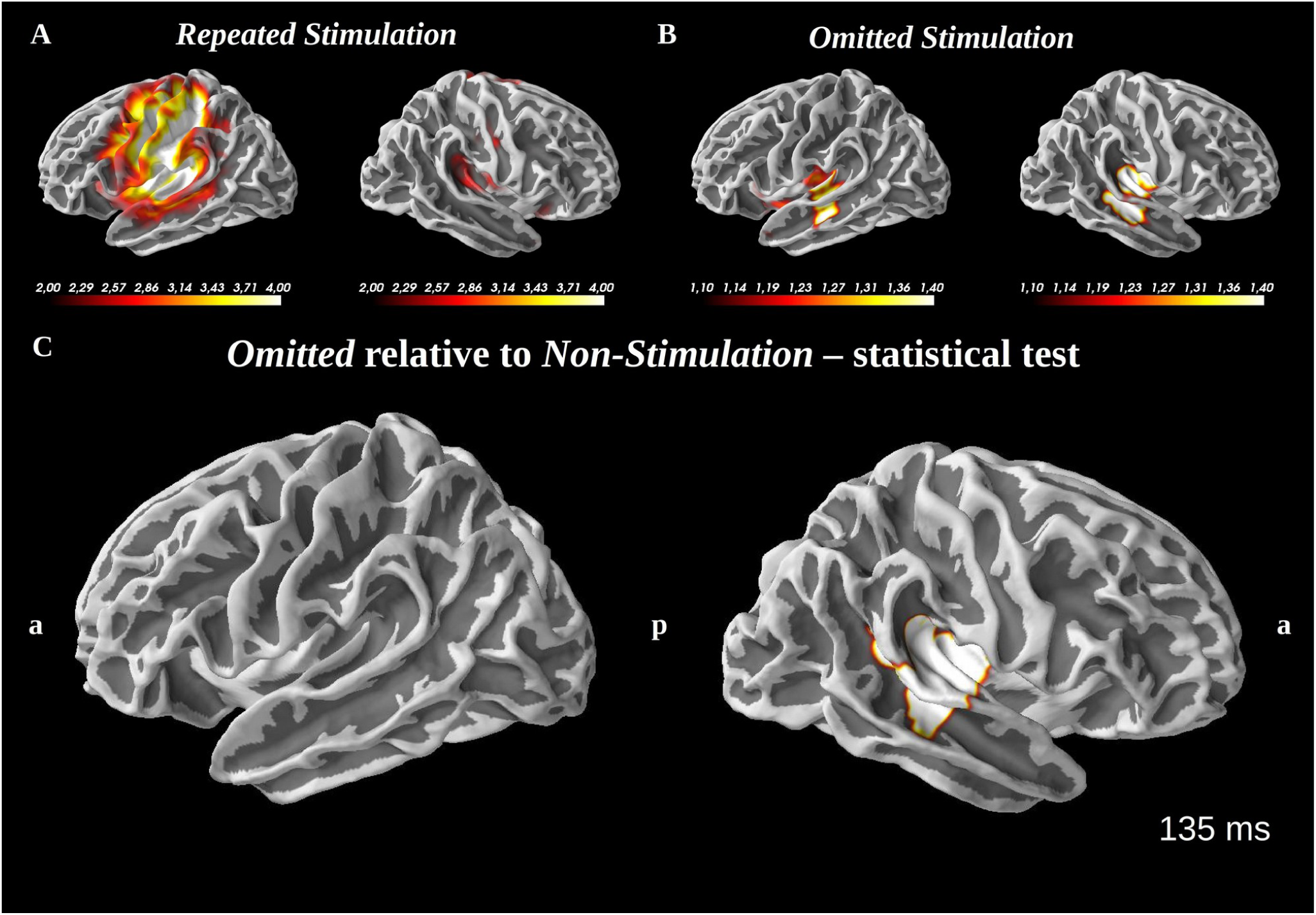
Timelocked responses following repeated and omitted stimulations. Grand averages (dSPM values) for *Repeated* and *Omitted Stimulation* and a statistical map based on cluster analysis for SII-component for *Omitted* versus *Non-Stimulation* after 135 ms**: A)** grand average source activity for *Repeated Stimulation.* This revealed bilateral activation of the SII and contralateral activation of SI. **B)** grand average source activity for *Omitted Stimulation.* This revealed bilateral activation of SII. **C)** *t*-maps cluster-thresholded at 0.025 overlaid on the fsaverage brain at 135 ms. The difference response is localized to the right superior temporal gyrus, posterior insula and SII. a=anterior, p=posterior.

#### Expectations during stimulations: *Repeated* versus *First Stimulation*

The influence from expectations on stimulations were explored by testing the differences between *Repeated* and *First Stimulation* for the two response components at 56 ms and 135 ms. The *p*-values for the biggest clusters were respectively: SI: *p =* 0.58; SII *p* = 0.21. Thus, the results showed no significant differences between *Repeated* and *First Stimulation* for any of these two response components.

#### Expectations in the absence of stimulation: *Omitted* versus *Non-Stimulation*

The influence from expectations during omissions were explored by testing the differences between *Omitted* and *Non-Stimulations* for the same two peaks, 56 ms and 135 ms. The *p-*values for the biggest clusters were respectively: SI: *p* = 0.58; SII: *p* = 0.0001. Thus, the results showed a significant difference between *Omitted* and *Non-Stimulation* for the second (but not the first) response component driven by more activity in ipsilateral SII for *Omitted* compared to *Non-Stimulation*.

### Induced responses

For induced responses, grand averages were calculated across all participants. Similar comparisons between conditions were made here as for the evoked responses, that is, comparing *Repeated* and *First Stimulation* on the one hand, and comparing *Omitted* and *Non-Stimulation* on the other. For both sets of analyses, we baselined the induced responses with the induced activity from the *No-Task* trials (segments of rest data before the task begun, but with the movie running).

#### The effect of stimulation

Investigating from stimulus onset till 1000 ms post-stimulus, we found the classical responses to tactile sensory stimulation (Salmelin et al., 1995; Salmelin and Hari, 1994), which include mu-alpha (∼12 Hz) and mu-beta (∼22 Hz) suppression from 150 ms to 500 ms and a mu-beta (∼18 Hz) rebound from 500 ms to 900 ms (Fig. 4). Furthermore, a theta synchronization (∼7 Hz) from −100 ms to 350 ms was found for both *Repeated* and *First Stimulation*, but with seemingly greater power for *Repeated* than for *First Stimulation* (Fig. 4B). For *Omitted* and *Non-Stimulation* these components were not clearly found (Fig. 4B).

**Fig. 4.**
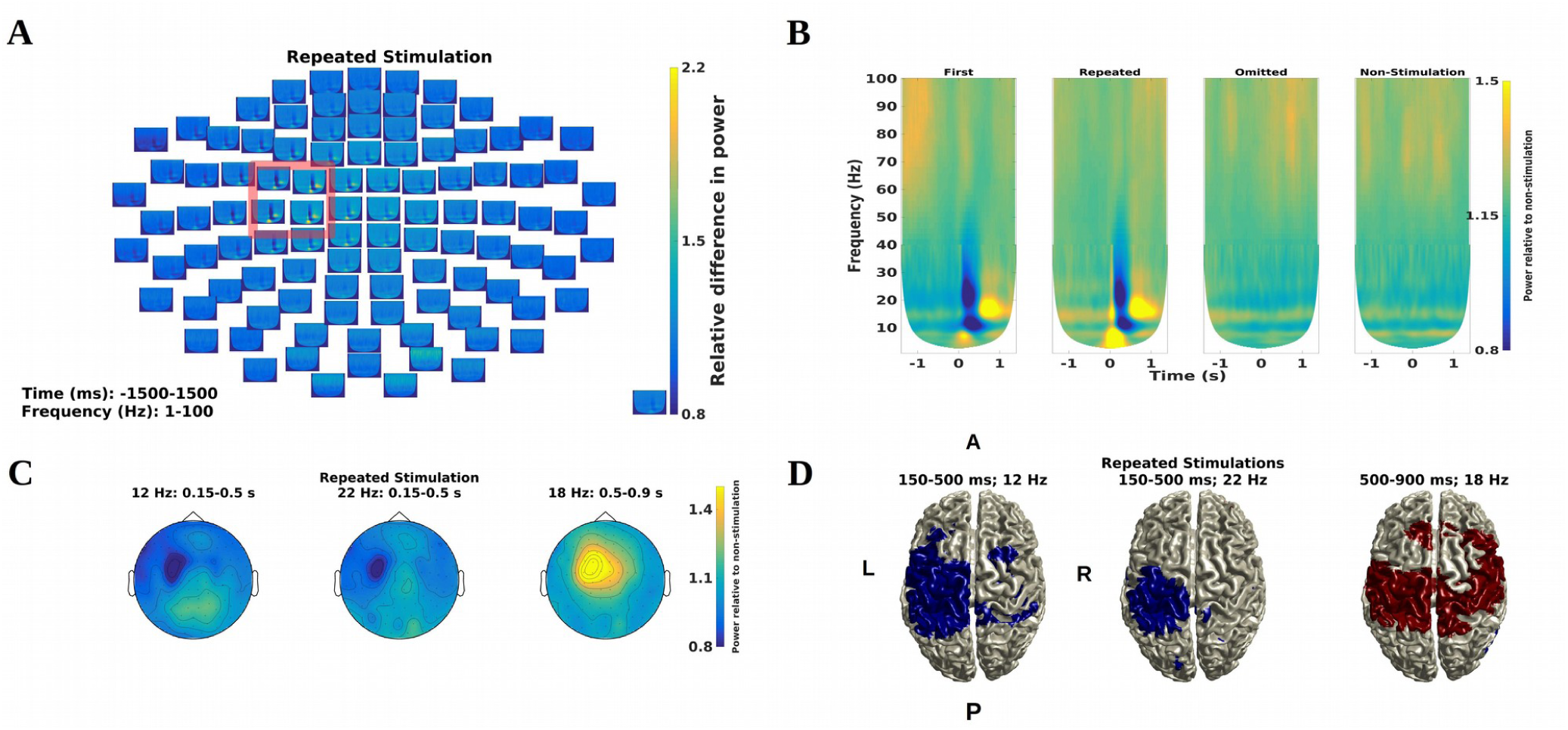
Induced responses during stimulation and during the absence of stimulation: Grand average time-frequency representations for gradiometer pairs. **A)** *Repeated Stimulation* channel plots. **B)** Mean of channels in the red square in **A)** for all conditions (Table 1): both the mu-alpha and mu-beta bands suppressions and the mu-beta rebound are seen for the stimulations, but not for the absence of stimulation. **C)** Topographical plots for *Repeated Stimulation,* showing a contralateral topography for all alpha and beta synchronizations and desynchronizations. **D)** Beamformer surface source reconstructions of the activity underlying the topographical plots, with activity significant at an alpha level of 0.05 (Red is greater than zero, blue is lesser than zero). For all plots, power is relative to the power for the *No-Task* trials recorded outside the experiment.

#### Expectations during stimulations: *Repeated* versus *First Stimulation*

A comparison between *Repeated* and *First Stimulation* revealed higher power for *Repeated* than for *First Stimulation* in the theta, beta and gamma bands (Fig. 5). The theta band increase was before and after the stimulation (∼7 Hz; from −100 ms to 350 ms). A beta band increase followed the stimulation directly (∼20 Hz; from 0 ms to 100 ms). It is not likely that these differences were related to phase-locked activity since no differences were found between *Repeated* and *First Stimulation* in the time-locked responses (Fig. 2AB). The gamma band increase was found in the pre-stimulus time period (−300 ms to 0 ms at ∼47 Hz). Finally, the beta band showed increased activity pre-stimulus (∼20 Hz; from −1300 ms to 0 ms). These four increases in synchronization were all identified in the biggest cluster (*p <* 0.001) when testing the differences between *Repeated* and *First Stimulation* (Fig. 5DE).

**Fig. 5.**
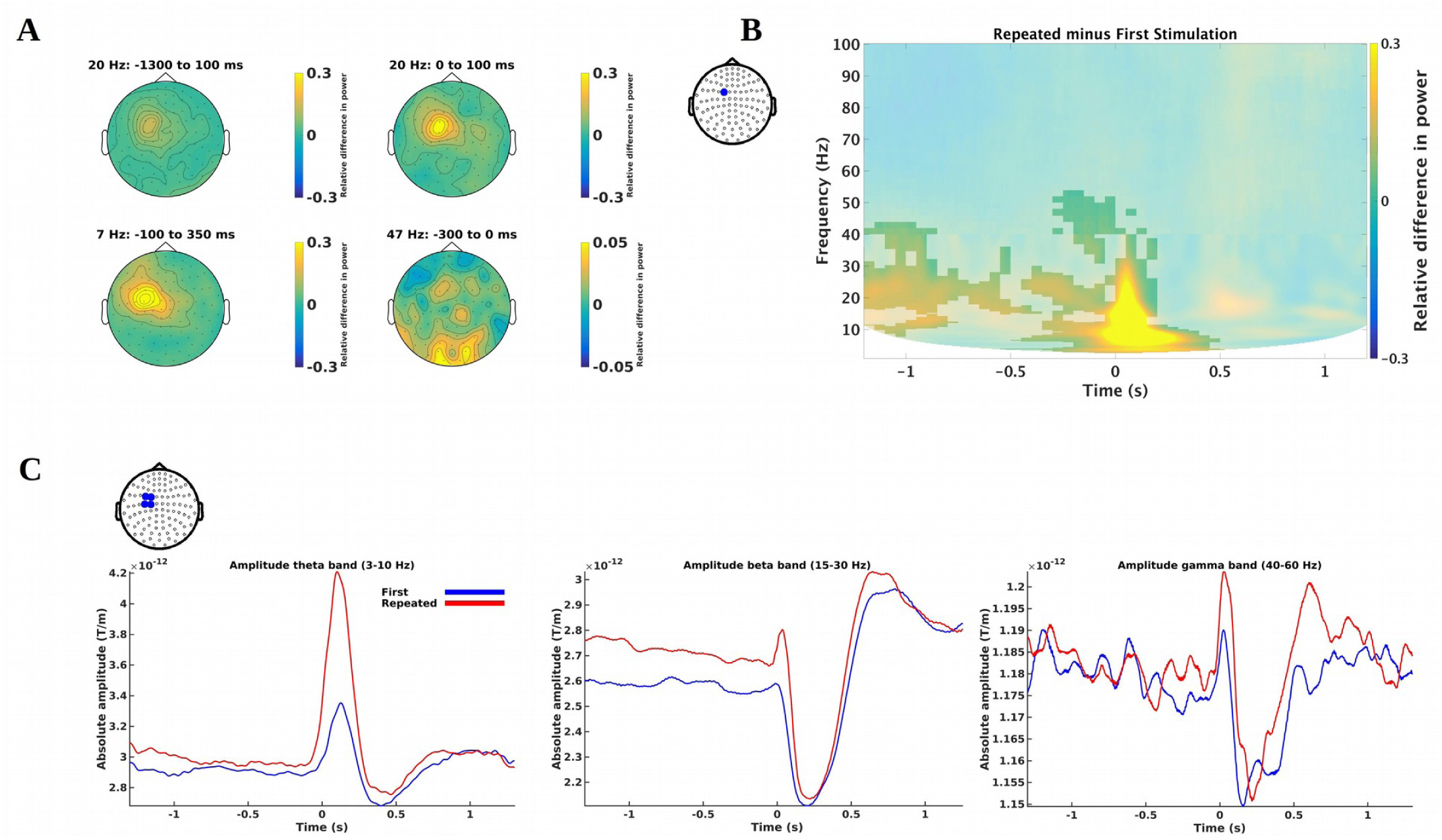
Differences in induced responses due to differences in expectations during stimulation: Differences between *Repeated and First Stimulation*. **A)** Topographical plots of theta, beta and gamma band differences **B)** Plot of a single channel showing the theta, beta and gamma band differences. Differences associated with a cluster with a *p*-value lower than an alpha of 0.025 are shown non-blurred. The position of the channel shown is indicated by the blue dot on the topography. **C)** Temporal spectral evolution plots of the theta, beta and gamma bands. These are based on an average of the four channels indicated by the blue dots on the topography.

Since it is potentially possible that the theta, beta and gamma band increases for *Repeated* relative to *First Stimulation* are simply due to refractory activity from the preceding stimulation, we examined this possibility by calculating the temporal spectral evolution of these frequency bands (Salmelin and Hari, 1994) (Fig. 5E). We demeaned the resulting time courses by taking the mean of the activity from −1300 ms to −500 ms to ensure that any differences found were not consequences of offset differences. We then tested using cluster statistics whether these peaks for *Repeated Stimulation* were significantly higher than the corresponding peaks for *First Stimulation*. The results showed that the increases in power for *Repeated* compared to *First Stimulation* (Fig. 5) were statistically significant for the theta, *p* < 0.001, and the beta bands, *p* = 0.0015, but not for the gamma band, *p* = 0.0575. The clusters, respectively, for the theta and the beta bands, extended from 19 ms to 232 ms and from 15 ms to 92 ms, closely matching the periods found in the induced responses. This indicates that the increases in induced responses found for *Repeated* relative to *First Stimulation* cannot be explained simply by increased activity due to the preceding stimulation.

#### Expectations in the absence of stimulation: *Omitted* versus *Non-Stimulation*

A comparison between *Omitted* and *Non-Stimulation* revealed lower power for *Omitted Stimulation* than for *Non-Stimulation* in the alpha band (Fig. 6). The decrease was found both before and after time point of the omitted stimulation (∼10 Hz; from −500 ms to 1000 ms). This decrease in synchronization was identified in the biggest cluster (*p <* 0.001) when testing the differences between *Non-Stimulation* and *Omitted Stimulation* (Fig. 6B).

**Fig. 6.**
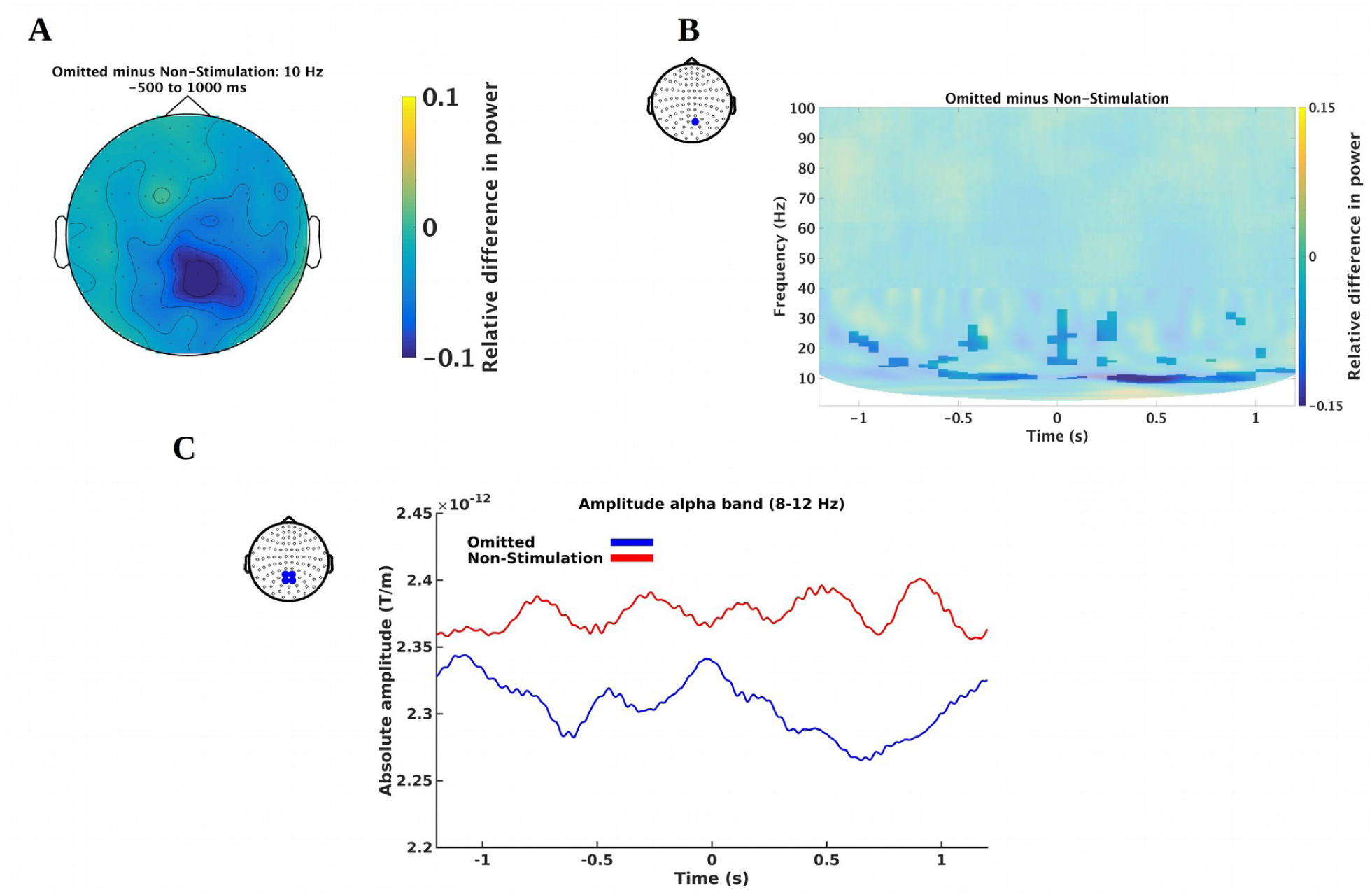
Differences in induced responses due to differences in expectations during during the absence of stimulation: Differences between *Omitted* and *Non-Stimulations*. **A)** Lower power in the alpha band (10 Hz) for *Omitted* relative to *Non-Stimulation*. **B)** Plot of a single channel showing an alpha band difference. Differences associated with a cluster with a *p*-value lower than an alpha of 0.025 are shown non-blurred. The position of the channel shown is indicated by the blue dot on the topography. **C)** Temporal spectral evolution plots of the alpha band. The lines are based on an average of the four channels indicated by the blue dots on the topography.

### Beamformer source reconstructions

The above results from the analysis of induced responses were followed up with source space analyses below. We performed beamformer source reconstructions for the alpha, beta, theta and gamma bands activity using the methods described in the methods section. Statistical tests for the *a priori* areas of interest (insula, thalamus, MCC, MFG, SI, IPC, IFG and cerebellum) were extracted below. For each of the areas the maximum unsigned value was extracted from each subject for that parcellation according to the AAL-atlas (Tzourio-Mazoyer et al., 2002). The statistical within-subject, between-conditions testing was done based on these values (Figs. 7-8 and Tables 2-3). No corrections for multiple comparisons were done on the beamformer reconstructions since the alpha level had already been controlled at α = 0.05 by the earlier permutation test (i.e. had there been no significant effect of the permutation test, no beamformer reconstructions would have been done). Note that the IFG in this study is defined as Brodmann Area 44, based on the coordinates supplied in Fardo et al. (2017).

**Table 2.**
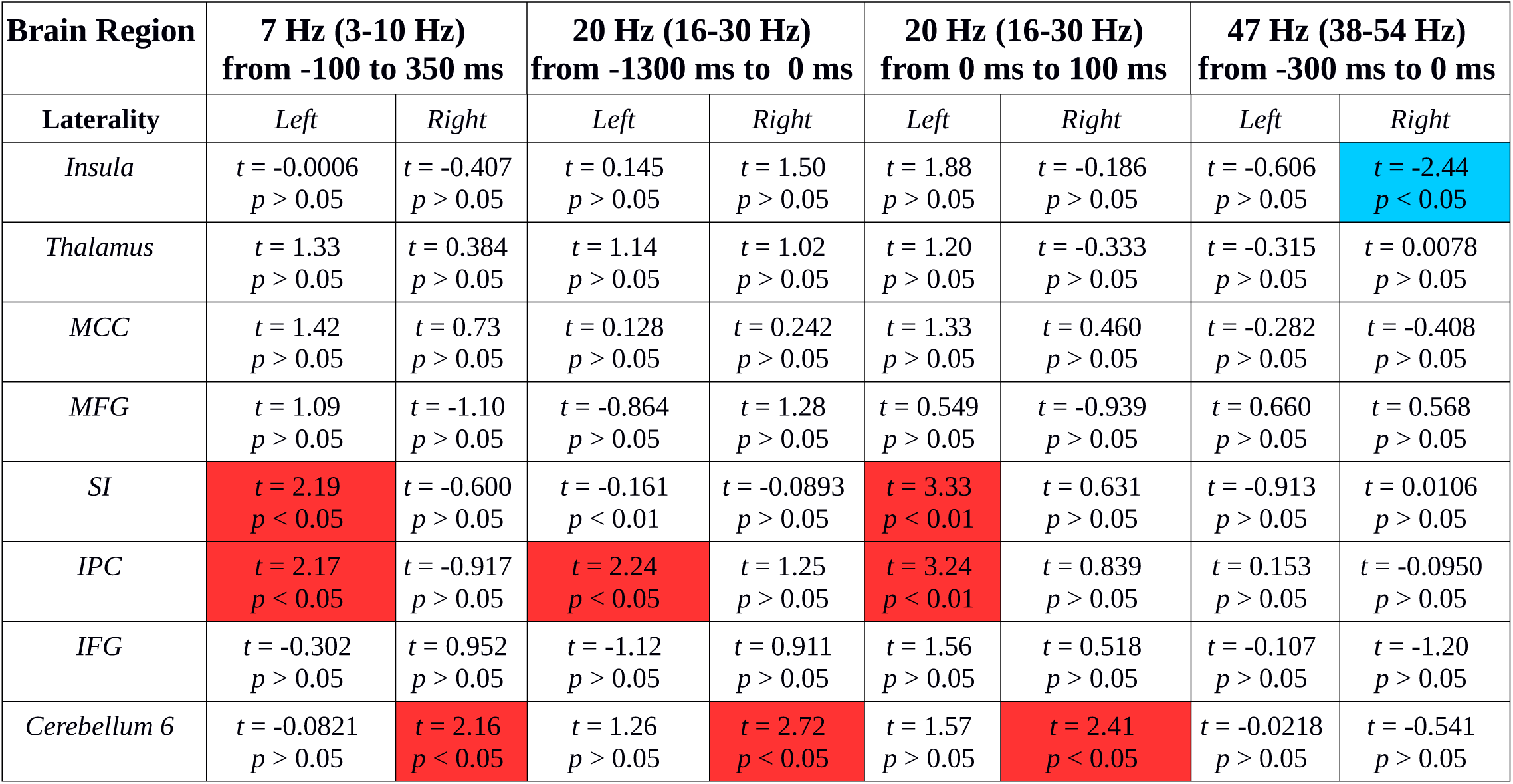
Tests of expectations during stimulation for the *a priori* areas: Statistical tests for stimulation main effects (*df* = 19) for areas reported by Allen et al. (2016), Fardo et al. (2017) and the cerebellum (Tesche and Karhu, 2000). **Red:** significantly greater activity for *Repeated* compared to *First Stimulation*. **Blue:** significantly greater activity for *Repeated* compared to *First Stimulation*. An alpha threshold of 0.05 is used.

| Brain Region | 7 Hz (3-10 Hz)<br>from -100 to 350 ms |  | 20 Hz (16-30 Hz)<br>from -1300 ms to 0 ms |  | 20 Hz (16-30 Hz)<br>from 0 ms to 100 ms |  | 47 Hz (38-54 Hz)<br>from -300 ms to 0 ms |  |
| --- | --- | --- | --- | --- | --- | --- | --- | --- |
| Laterality | Left | Right | Left | Right | Left | Right | Left | Right |
| *Insula* | $t = -0.0006$<br>$p > 0.05$ | $t = -0.407$<br>$p > 0.05$ | $t = 0.145$<br>$p > 0.05$ | $t = 1.50$<br>$p > 0.05$ | $t = 1.88$<br>$p > 0.05$ | $t = -0.186$<br>$p > 0.05$ | $t = -0.606$<br>$p > 0.05$ | $t = -2.44$<br>$p < 0.05$ |
| *Thalamus* | $t = 1.33$<br>$p > 0.05$ | $t = 0.384$<br>$p > 0.05$ | $t = 1.14$<br>$p > 0.05$ | $t = 1.02$<br>$p > 0.05$ | $t = 1.20$<br>$p > 0.05$ | $t = -0.333$<br>$p > 0.05$ | $t = -0.315$<br>$p > 0.05$ | $t = 0.0078$<br>$p > 0.05$ |
| *MCC* | $t = 1.42$<br>$p > 0.05$ | $t = 0.73$<br>$p > 0.05$ | $t = 0.128$<br>$p > 0.05$ | $t = 0.242$<br>$p > 0.05$ | $t = 1.33$<br>$p > 0.05$ | $t = 0.460$<br>$p > 0.05$ | $t = -0.282$<br>$p > 0.05$ | $t = -0.408$<br>$p > 0.05$ |
| *MFG* | $t = 1.09$<br>$p > 0.05$ | $t = -1.10$<br>$p > 0.05$ | $t = -0.864$<br>$p > 0.05$ | $t = 1.28$<br>$p > 0.05$ | $t = 0.549$<br>$p > 0.05$ | $t = -0.939$<br>$p > 0.05$ | $t = 0.660$<br>$p > 0.05$ | $t = 0.568$<br>$p > 0.05$ |
| *SI* | $t = 2.19$<br>$p < 0.05$ | $t = -0.600$<br>$p > 0.05$ | $t = -0.161$<br>$p < 0.01$ | $t = -0.0893$<br>$p > 0.05$ | $t = 3.33$<br>$p < 0.01$ | $t = 0.631$<br>$p > 0.05$ | $t = -0.913$<br>$p > 0.05$ | $t = 0.0106$<br>$p > 0.05$ |
| *IPC* | $t = 2.17$<br>$p < 0.05$ | $t = -0.917$<br>$p > 0.05$ | $t = 2.24$<br>$p < 0.05$ | $t = 1.25$<br>$p > 0.05$ | $t = 3.24$<br>$p < 0.01$ | $t = 0.839$<br>$p > 0.05$ | $t = 0.153$<br>$p > 0.05$ | $t = -0.0950$<br>$p > 0.05$ |
| *IFG* | $t = -0.302$<br>$p > 0.05$ | $t = 0.952$<br>$p > 0.05$ | $t = -1.12$<br>$p > 0.05$ | $t = 0.911$<br>$p > 0.05$ | $t = 1.56$<br>$p > 0.05$ | $t = 0.518$<br>$p > 0.05$ | $t = -0.107$<br>$p > 0.05$ | $t = -1.20$<br>$p > 0.05$ |
| *Cerebellum 6* | $t = -0.0821$<br>$p > 0.05$ | $t = 2.16$<br>$p < 0.05$ | $t = 1.26$<br>$p > 0.05$ | $t = 2.72$<br>$p < 0.05$ | $t = 1.57$<br>$p > 0.05$ | $t = 2.41$<br>$p < 0.05$ | $t = -0.0218$<br>$p > 0.05$ | $t = -0.541$<br>$p > 0.05$ |

**Fig. 7.**
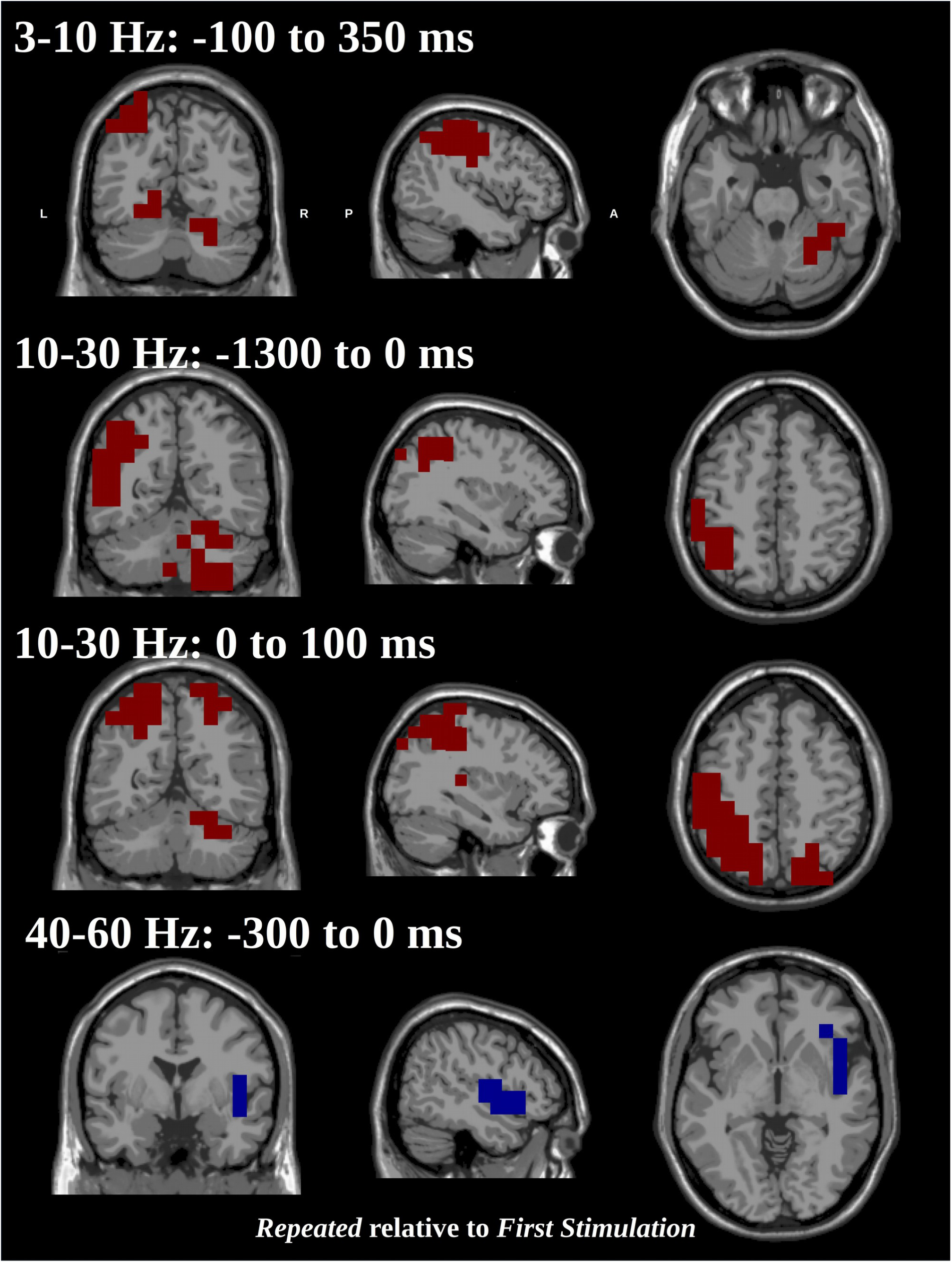
Statistical *t-*maps of areas showing significant power differences based on beamformer reconstructions of stimulations: Both the theta and beta bands showed more activity for *Repeated* than for *First Stimulations* in the right-lateralized cerebellum and in the left inferior parietal cortex. SI, however, only showed increased activity at the times around stimulation. The gamma band showed greater activity for *First* than for *Repeated Stimulation* in the right insula **Red:** significantly greater activity for *Repeated* compared to *First Stimulation.* **Blue:** significantly greater activity for *Repeated* compared to *First Stimulation*. An alpha threshold of 0.05 is used. Tests are run across parcellations based on the AAL-atlas. Axis keys: L=Left, R=Right, A=Anterior, P=Posterior. The slices viewed are respectively: *x*_*1*_ *=* −44 mm, *y*_*1*_ = −65 mm, *z*_*1*_ = −26 mm; *x*_*2*_ *=* −35 mm, *y*_*2*_ = −57 mm, *z*_*2*_ = 48 mm; *x*_*3*_ *= -*35 mm, *y*_*3*_ = −57 mm, *z*_*3*_ = 48 mm; *x*_*4*_ *=* 48 mm, *y*_*4*_ = −1 mm, *z*_*4*_ = −4 mm, all in MNI space.

#### Beamformer reconstructed activity

The activity related to the mu-alpha band (12 Hz, from 150 ms to 500 ms) and to the mu-beta band (22 Hz, from 150 ms to 500 ms; and from 500 ms to 900 ms, 18 Hz) were localized to the contralateral somatosensory and motor cortices (see Fig. 4D).

These source localizations replicate what has been reported in the literature for induced responses of tactile stimulation before (Cheyne, 2013) and thus serve as a sanity check that our stimulation worked as intended. Since no differences were found between *Repeated* and *First Stimulation* in the induced responses for these bands, no statistical comparisons were made. Note that no parcellation was used for these alpha and beta bands analyses, since these only served as sanity checks.

#### Expectations during stimulations: *Repeated* versus *First Stimulation*

The role of expectations during stimulation was tested by contrasting *Repeated* against *First Stimulation.* All three bands, theta, beta and gamma, showed greater power over contralateral SI and IPC for *Repeated* as compared to *First Stimulations* (Fig. 7). The SI activity was however absent during the pre-stimulus period, −1300 to 0 ms. Also for *Repeated* contrasted to *First Stimulations*, the theta and beta bands showed greater power over the right cerebellum (Fig. 7). The theta and beta bands revealed very similar patterns of activations. Finally, for *First* contrasted against *Repeated Stimulations* greater activation in the gamma band was found in the right insula (Fig. 7).

#### Expectations in the absence of stimulation: *Omitted* versus *Non-Stimulation*

The role of expectations during absence of stimulation was tested by contrasting *Omitted* to *Non-Stimulation*. For this contrast, the alpha band revealed greater synchronization in the right cerebellum (Fig. 8) for *Non-Stimulation* compared to *Omitted*. See Table 3 for a summary of all the a priori areas.

**Table 3.** Test of expectations during absence of stimulation for the *a priori* areas: Statistical tests for stimulation main effects (*df* = 19) for areas reported by Allen et al. (2016), Fardo et al. (2017) and the cerebellum(Fardo et al., 2017) (Tesche and Karhu, 2000). **Blue:** significantly greater activity for *Non-Stimulation* compared to *Omitted*. An alpha threshold of 0.05 is used.

| Brain Region | 10 Hz from 0 ms to 1110 ms |  |
| --- | --- | --- |
| Laterality | Left | Right |
| *Insula* | $t = -0.513$<br>$p > 0.05$ | $t = 0.193$<br>$p > 0.05$ |
| *Thalamus* | $t = 0.0581$<br>$p > 0.05$ | $t = -2.01$<br>$p > 0.05$ |
| *MCC* | $t = -0.945$<br>$p > 0.05$ | $t = -1.02$<br>$p > 0.05$ |
| *MFG* | $t = 1.62$<br>$p > 0.05$ | $t = 1.36$<br>$p > 0.05$ |
| *SI* | $t = 0.260$<br>$p > 0.05$ | $t = -1.40$<br>$p > 0.05$ |
| *IPC* | $t = -0.210$<br>$p > 0.05$ | $t = -1.87$<br>$p > 0.05$ |
| *IFG* | $t = 0.768$<br>$p > 0.05$ | $t = -0.256$<br>$p > 0.05$ |
| *Cerebellum 6* | $t = 0.921$<br>$p > 0.05$ | $t = -2.42$<br>$p < 0.05$ |

**Fig. 8.**
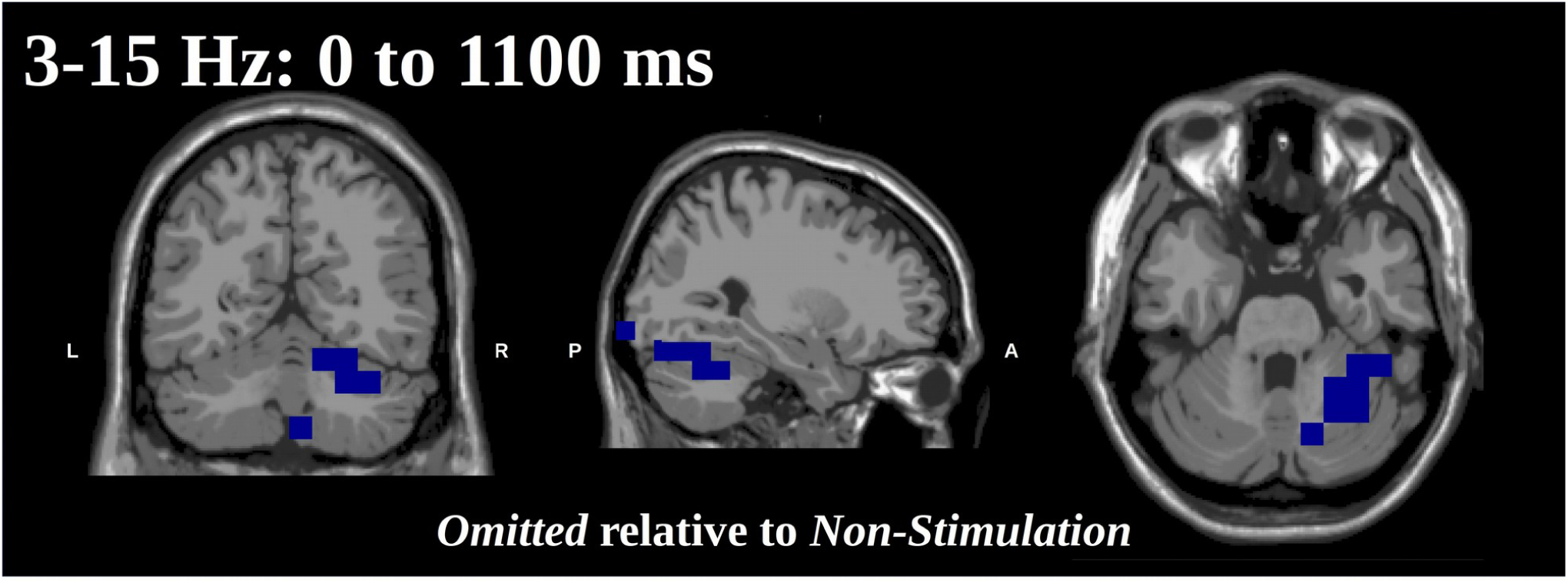
Statistical *t-*maps of areas showing significant power differences based on beamformer reconstructions of stimulations: The alpha band showed less activity for *Omitted* than for *Non-Stimulation* in the right-lateralized cerebellum. **Blue:** significantly greater activity for *Non-Stimulation* compared to *Omitted.* An alpha threshold of 0.05 is used. Tests are run across parcellations based on the AAL-atlas. Axis keys: L=Left, R=Right, A=Anterior, P=Posterior. The slices viewed are: *x*_*1*_ *=* 28 mm, *y*_*1*_ = −59 mm, *z*_*1*_ = −29 mm in MNI space.

## Discussion

In this study, we explored how expectations of somatosensory stimulations are established and expressed in neural response patterns as measured with MEG. Using experimental conditions with actual stimulation and omitted stimulations, both of which were either expected or unexpected, we aimed at elucidating the expression of expectations both during stimulations and during omissions.

### A precisely time- and phase-locked manifestation of expectation

As an initial data quality control, we observed the “classical” responses subsequent to all our actual stimulation events, both in terms of time-locked responses (Fig. 2AB) and induced responses in the time-frequency domain (Fig. 4). The time-locked responses hence displayed the typical somatosensory SI (∼60 ms) and SII (∼135 ms) components (Hari and Forss, 1999); and the induced responses displayed the expected mu-alpha and mu-beta patterns with desynchronization at ∼12 and ∼22 Hz from 150 ms to 500 ms and a beta rebound at ∼18 Hz from 500 ms to 900 ms (Fig. 4CD) (Salmelin et al., 1995; Salmelin and Hari, 1994).

Our study also provide new findings: When a stimulation is expected but omitted, there is a time-locked SII and right insular response occurring at ∼135 ms subsequent to the expected stimulation (Fig. 3). These results revealed a precisely time-and phase-locked response following the expected onset of an *Omitted Stimulation*, despite an inter-stimulus interval of 3000 ms (Fig. 2C), and thus provide novel evidence of the capability of the brain for doing very precise time-keeping of expected events across extended intervals. This finding is particularly surprising, considering related results from the auditory modality, where time-locked responses to omitted stimulations have only been demonstrated when the inter-stimulus interval is twenty times shorter (less than 150 ms) (Yabe et al., 1997). The difference in time-keeping across inter-stimulus intervals between the auditory and somatosensory systems might be due to the differences between the quality of auditory stimuli (such as speech and environmental sounds), which typically are brief and abrupt, and the quality of tactile stimuli, which are often comparatively slow and prolonged. Note however that a recent study of Naeije et al. (2018) did not find any evidence for timelocked activity related to omitted somatosensory stimulations when the inter-stimulus interval was 500 ms event though their source reconstructions also localized the source to SII. The disagreement between their results and our present results indicate that longer inter-stimulus interval than 500 ms 3000 ms may be necessary to generate SII responses of a sufficient amplitude for identification.

### Theta, beta, and gamma increases after expected stimulations

Our study also shows that when comparing *Repeated Stimulation* to *First Stimulation,* and hence isolating the influence of expectations during stimulations, there is an increase in theta band synchronization (∼7 Hz, from −150 ms to 350 ms) and beta band synchronization (∼20 Hz, from 0 ms to 100 ms) associated with expectation (Fig. 5). For these two increases in synchronization, beamforming revealed increased activity in *Repeated* relative to *First Stimulation* in somatosensory, parietal and cerebellar sources (Fig. 7 & Table 2). Leading up to the stimulation, we also found increased beta (∼20 Hz, from −1300 ms to 0 ms) and gamma band synchronization (∼47 Hz, from −300 ms to 0 ms) (Fig. 5). For the increase in beta band synchronization before stimulation, beamforming revealed parietal and cerebellar sources, but no somatosensory sources. This indicates that stimulations are followed by refractory activity in the inferior parietal cortex and the cerebellum and that these synchronizations result in the increase in synchronization of SI theta and beta band activity (Allen et al., 2016). A similar role of inferior parietal cortex has also been reported by Fardo et al. (2017), where results from localizations of event-related fields from a tactile oddball indicated that inferior parietal cortex was involved in updating expectations.

One could possibly argue that the differences in theta and beta between *Repeated and First Stimulation* (Figs. 5 & 7 & Table 2) could simply be interpreted as gating activity, (Arnfred et al., 2001; Swerdlow et al., 1992), attenuating the magnitude of the subsequent, and expected stimulations. If this was the case, the observed increase in power would reflect increased inhibition of the processing of such expected stimulations. Two things however speak against such an interpretation. First, no habituation effects were observed on the evoked fields (Fig. 2), while habituation has been reported in studies with shorter inter-stimulus intervals (< 2 s) (Cheng et al., 2017; Hsiao et al., 2013). Second, conversely to what was found here, the influence from gating on beta oscillations have been shown to show *higher* synchronization for the first than for the second stimulation when stimulations are presented in pairs (Hsiao et al., 2013). The increase in theta and beta power between *Repeated* and *First Stimulation* in our results hence appears to be a true manifestation of expectation rather than a gating phenomenon.

Another interpretation of the observed increase in beta power between *Repeated Stimulations* and *First Stimulation* is that the beta band would be signalling the status quo as according to Engel and Fries (2010). An intuitive conception of what the status quo amounts to is exemplified by idling rhythms, e.g. the mu-rhythm over central sensors and the alpha rhythm over posterior sensors, when the subject is at rest (Niedermeyer and Silva, 2005). Engel and Fries, however, extended the idea for status quo from an idling rhythm (Pfurtscheller et al., 1996) to also include the perceptual set, the sensory expectations, where they hypothesize that maintenance of the sensory expectation would cause increased synchronization in the beta band. Such an increase may hence be what we see in the beta band change at stimulation, 0 to 100 ms, (Figs. 5 & 7) from *First Stimulation*, where no sensory expectation is yet established, to *Repeated Stimulation*, where the sensory expectation is established and needs to be maintained, which the present results would indicate would be done by inferior parietal cortex and cerebellum (−1300 ms to 0 ms) (Fig. 7 & Table 2).

The present results may seem to be in opposition to earlier results where expected and attended tactile stimuli were accompanied by desynchronizations in the beta band (van Ede et al., 2011, 2010). These previous experiments, however included an *active* task for the participants, contrary to the current experimental *passive* protocol. This means that that in those previous studies, the tactile stimulations must be processed attentively for the research participant to perform the task at an acceptable level. In the current study, no tasks were involved, and hence no such attentive or otherwise active processing was necessary. Rather, subjects were engaged with looking at a documentary movie and if anything directing their attention away from the tactile stimulations.

The gamma differences were wholly related to less activity for *Repeated* relative to *First Stimulation* activity in the right insula, an area which has been reported to be related to anticipation for the consequences of touch (Lovero et al., 2009) and coordinating activity related to prediction errors regarding upcoming stimulation (Allen et al., 2016). The gamma-band activity likely serves the purpose of updating the internal state in the network, with right insula signalling that a new chain of stimulations has begun (Allen et al., 2016) (Fig. 7 & Table 2).

Neither the thalamus, the MCC nor the frontal gyri all included among the *a priori* areas - showed any differences between the conditions. However, both the thalamus and the MCC are in areas where MEG shows little sensitivity (Hillebrand and Barnes, 2002), and hence the present study might be underpowered to find them. Also, in the study of Fardo et al. (2017) the IFG showed effects relative to attention and not directly to expectation.

Between *Non-Stimulations and Omitted Stimulations –* reflecting the role of expectations in the absence of stimulations *–* we found a difference in the alpha band (∼10 Hz, from 0 ms to 1100 ms) (Fig. 6). This comparison revealed lower power in the right cerebellum for *Omitted Stimulations* compared to *Non-Stimulations* The activity difference does not emerge as related to the expected onset of the *Omitted Stimulation*, but rather as a continuing desynchronization, as compared to *Non-Stimulation* (Fig. 6D). At this moment, it is not entirely clear what this represents.

### Role of cerebellum and parietal cortex – maintaining the status quo?

Our results also showed power differences in the right cerebellum for the theta and beta bands, with more activation for *Repeated* than *First Stimulation* (Figs. 7-8 and Tables 2-3). The refractory activity in the beta band after a stimulation (Fig. 5) was found in left inferior parietal cortex and the right cerebellum.

A tentative interpretation of this is that during tactile stimulation, *First Stimulation*, activates continuing cerebellar and parietal responses and that each new stimulation, *Repeated Stimulation*, is accompanied by stronger SI activation at stimulation due to these continuing cerebellar and parietal activities. In this sense, the refractory cerebellar activity and inferior parietal cortex in the beta band may be responsible for maintaining the status quo (Engel and Fries, 2010). To strengthen this interpretation, it would however be necessary to find dissociative evidence, such as cases where there is no beta band peak at the time of stimulation (as in Fig. 5E) even though the stimulation is a repetition. From the earlier literature (Tesche and Karhu, 2000), it has been suggested that the cerebellar activity has a refractory period of 2-4 s. Future studies could therefore aim at varying the inter-stimulus interval and including intervals beyond this refractory period. Given the refractory period of 2-4 s for cerebellar activity, it would furthermore be interesting to investigate how dependent the time-locked effect is on the duration between stimulations, within and outside the 2-4 second time window. Indeed, the results of Naeije et al. (2018) indicate that there might also be a lower limit on when this effect can be detected, as indicated by the absence of a significant effect for omitted stimuli when the inter-stimulus interval was 500 ms.

One thing that one must always consider in MEG studies is how much credibility one is willing to assign to subcortical localizations. The cerebellum gains credibility by having been detected in earlier studies (Tesche and Karhu, 2000) and also from the theoretical knowledge that cerebellum is activated ipsilaterally to stimulation, as was also found in the current study (Fig. 10). Also more and more studies are surfacing for MEG being sensitive to deep sources (Attal et al., 2012; Coffey et al., 2016; Garrido et al., 2015; Tenney et al., 2013; Tesche and Karhu, 1997),. For future work, it would be of great value to have more detailed models for the cerebellum such that the orientations and positions of potential sources can be modelled with greater accuracy and thus subsequently raise our belief in subcortical localizations.

## Conclusions

This study aimed at elucidating the expression of expectations both during actual tactile stimulations and during omitted stimulations. The results provide new insights into how the brain updates and maintains the expectations towards sensory touch. We show that neural processing of omissions occurs in a precisely time-locked manner, and that it is generated by posterior insula and SII for the time-locked responses. This indicates that the brain keeps a very precise timing of when events are expected to happen even across intervals of 3000 ms, well beyond what has been earlier reported in the literature. We also show that gamma band activity is involved in updating the brain about new stimulations. In this way the insula plays a dual role, showing activity that correlates both with omitted stimulations and with the first stimulation of new chains of stimulation.

Refractory beta band activity was found in the cerebellum and the inferior parietal cortex after a stimulation. Extra involvement of SI when stimulations were repeated was also found. This may be interpreted as the beta band signalling the status quo – that a predictable sequence of stimulations is expected. The theta band also showed cerebellar, inferior parietal cortex and SI activity for repeated stimulations relative to new stimulations.

## Acknowledgements

The authors wish to thank Robert Oostenveld for valuable comments on an earlier version of the design and to thank Veikko Jousmäki for valuable comments on the interpretation of the results.

Data for this study was collected at NatMEG (www.natmeg.se), the National infrastructure for Magnetoencephalography, Karolinska Institutet, Sweden. The NatMEG facility is supported by Knut & Alice Wallenberg (KAW2011.0207). The study, and Lau Møller Andersen, was funded by Knut & Alice Wallenberg Foundation (KAW2014.0102).

